# Integrated time course omics analysis distinguishes immediate therapeutic response from acquired resistance

**DOI:** 10.1101/136564

**Authors:** Genevieve Stein-O’Brien, Luciane T Kagohara, Sijia Li, Manjusha Thakar, Ruchira Ranaweera, Hiroyuki Ozawa, Haixia Cheng, Michael Considine, Sandra Schmitz, Alexander V Favorov, Ludmila V Danilova, Joseph A Califano, Evgeny Izumchenko, Daria A Gaykalova, Christine H Chung, Elana J Fertig

## Abstract

**BACKGROUND:** Targeted therapies specifically act by blocking the activity of proteins that are encoded by genes critical for tumorigenesis. However, most cancers acquire resistance and long-term disease remission is rarely observed. Understanding the time course of molecular changes responsible for the development of acquired resistance could enable optimization of patients’ treatment options. Clinically, acquired therapeutic resistance can only be studied at a single time point in resistant tumors. To determine the dynamics of these molecular changes, we obtained high throughput omics data weekly during the development of cetuximab resistance in a head and neck cancer *in vitro* model.

**RESULTS:** An unsupervised algorithm, CoGAPS, was used to quantify the evolving transcriptional and epigenetic changes. Applying a PatternMarker statistic to the results from CoGAPS enabled novel heatmap-based visualization of the dynamics in these time course omics data. We demonstrate that transcriptional changes result from immediate therapeutic response or resistance, whereas epigenetic alterations only occur with resistance. Integrated analysis demonstrates delayed onset of changes in DNA methylation relative to transcription, suggesting that resistance is stabilized epigenetically.

**CONCLUSIONS:** Genes with epigenetic alterations associated with resistance that have concordant expression changes are hypothesized to stabilize resistance. These genes include *FGFR1,* which was associated with EGFR inhibitor resistance previously. Thus, integrated omics analysis distinguishes the timing of molecular drivers of resistance. Our findings provide a relevant towards better understanding of the time course progression of changes resulting in acquired resistance to targeted therapies. This is an important contribution to the development of alternative treatment strategies that would introduce new drugs before the resistant phenotype develops.

## BACKGROUND

Recent advances to identification of gene regulation in cancer has enabled the selection of targeted therapies to inhibit specific regulators of oncogenic signaling pathways essential for tumor development and maintenance [1]. These therapies prolong survival but are not curative, since most patients develop acquired resistance within the first few years of treatment [2]. Although a wide variety of molecular alterations that confer resistance to the treatment have been described, the mechanisms and timing of their evolution are still poorly characterized [3,4]. As serial biopsies along the treatment period are impractical due to the invasiveness and high costs of the procedure, the molecular alterations associated with acquired resistance are only known when resistance has already developed, and little is known about which changes occur at earlier or later time points in the course of the targeted therapy. The lack of adequate *in vitro* and *in vivo* time course datasets makes it challenging to delineate the two predominant hypotheses for how therapeutic resistance develops: (1) the presence of small populations of resistant cells that will survive the treatment and repopulate the tumor; or (2) the development of *de novo* resistance mechanisms by the tumor cells [4,5]. Characterization of the dynamics of genomic alterations induced during acquired resistance can identify targetable oncogenic drivers and determine the best time point to introduce alternative therapeutic strategies to avoid resistance establishment [6].

Epidermal Growth Factor Receptor (EGFR) inhibitors represent a common class of targeted therapeutics. Cetuximab, a monoclonal antibody against EGFR, is FDA approved for the treatment of metastatic colorectal cancer and head and neck squamous cell carcinoma (HNSCC) [7]. As with other targeted therapies, stable response is not observed for a long period and virtually all patients invariably develop acquired resistance [8]. Recent advances in the establishment of *in vitro* models of acquired cetuximab resistance [9] provide a unique opportunity to study the time course of genetic events resulting in acquired resistance. Cell lines chronically exposed to the targeted agent develop resistance and can be sequentially collected during the course of treatment to evaluate the progressive molecular changes. Previous studies to assess the mechanisms of acquired cetuximab resistance have been limited to comparing the genomic profile of the parental intrinsic sensitive cell line to stable clones with acquired resistance [9–11]. Therefore, these studies fail to capture the dynamics of acquired molecular alterations during the evolution of therapeutic resistance.

The development of *in vitro* time course data to determine the molecular drivers of therapeutic resistance is crucial. These experimental systems have the further advantage that time course data can also be generated for untreated controls, enabling the distinction of the molecular mechanisms associated with acquired resistance from those that would occur due to the long term culturing over the time period that resistance develops.

Along with novel time course datasets, inferring the specific and targetable signaling changes that drive therapeutic resistance also requires new bioinformatics pipelines to analyze and visualize these data. The bioinformatics pipelines must integrate genetic, epigenetic, and transcriptional changes from multiple-high throughput platforms to infer the complex gene regulatory mechanisms that are responsible for acquired resistance. Current supervised bioinformatics algorithms that find time course patterns in genomic data adjust linear models to correlate molecular profiles with known temporal patterns [12–15]. Many unknown variables such as culture conditions, immediate response to cetuximab, and adaptive changes may have confounding effects on known covariates of therapeutic response such as growth rates, colony size, or apoptosis rates. Unsupervised bioinformatics algorithms learn the dynamics directly from the high-throughput data, and therefore do not require *a priori* knowledge of the complex dynamics associated with therapeutic response. Some unsupervised algorithms [16–21] seek breaking points of coherent, regulatory relationships to infer the dynamics along pathways. Many of these algorithms trace individual phenotypes or individual genomics platforms. Their ability to determine drivers of gene expression associated with acquired resistance from time course data in multiple experimental conditions and multiple genomics data modalities is emerging [22]. Further extensions are needed to contrast the dynamics of signaling response to therapy to the dynamics of control conditions to distinguish the specific molecular processes that are unique to resistance. Matrix factorization algorithms are unsupervised, and can distinguish the relative molecular changes in each experimental condition over time without requiring prior knowledge of gene regulation. We have found that Bayesian, non-negative matrix factorization algorithms such as Coordinate Gene Activity in Pattern Sets (CoGAPS) [23] can extend beyond clustering to robustly quantify the dynamics and infer the gene regulatory networks directly from the input time course data [24]. The CoGAPS error model can also be modified to enable data-driven inference in distinct molecular platforms for inference of epigenetic regulation of gene expression [25].

In this study, we developed a new bioinformatics analysis pipeline for integrated analysis of gene expression and DNA methylation changes that occur during the time course progression of resistance to targeted therapies using CoGAPS. Genes uniquely associated with these changes were selected using a PatternMarker statistics [26] to enable novel visualization of molecular alterations dynamics inferred with CoGAPS. In order to benchmark our new bioinformatics pipeline, we used an *in vitro* HNSCC cell line model to induce resistance and measure the molecular changes using high throughput assays while the resistant phenotype developed. Gene expression and DNA methylation changes were screened weekly while acquired cetuximab resistance was induced in SCC25 cell line (intrinsic sensitive to cetuximab) and compared to the status of the untreated controls at the same culturing time point. CoGAPS [26] inferred specific patterns of expression and DNA methylation that are associated with the gradual establishment of acquired cetuximab resistance. The onset of methylation changes associated with resistance is temporally delayed relative to expression changes suggesting that epigenetic alterations stabilize the transcriptional changes relevant to the resistant phenotype. This analysis found anti-correlated changes between DNA methylation and gene expression in *FGFR1* during acquired therapeutic resistance. Up-regulation of *FGFR1* has previously been associated as a mechanism of acquired cetuximab resistance in HNSCC patients [27–29]. The identification of a canonical marker of resistance to EGFR inhibitors in this present study corroborates the efficacy of our experimental model and analytical algorithm to identify mechanisms of resistance. To our knowledge, this is the first demonstration of the anti-correlation between FGFR1 methylation and expression suggestive of its epigenetic regulation in acquired resistance to cetuximab. Thus, this pipeline can identify mechanisms of gene regulation in acquired resistance from high-throughput, multi-platform time course data. The resulting bioinformatics pipeline is poised to infer the dynamics of acquired resistance from emerging time course data with other cancer types and therapeutics.

## METHODS

### Cell lines and materials

SCC25 cells were purchased from American Type Culture Collection (ATCC). Cells were cultured in Dulbecco’s Modified Eagle’s medium and Ham’s F12 medium supplemented with 400ng/mL hydrocortisone and 10% fetal bovine serum and incubated at 37°C and 5% carbon dioxide. The parental cell line SCC25 and the late cetuximab and PBS generation 10 were authenticated using short tandem repeat (STR) analysis kit PowerPlex16HS (Promega, Madison, WI) through the Johns Hopkins University Genetic Resources Core Facility. Cetuximab (Lilly, Indianapolis, IN) was purchased from the Johns Hopkins Pharmacy.

### Induction of cetuximab resistance and time course sample collection

The HNSCC cell line SCC25 (intrinsically sensitive to cetuximab) was treated with 100nM cetuximab every three days for 11 weeks (generations G1 to G11). On the eighth day, cells were harvested. Sixty thousand cells were replated for another week of treatment with cetuximab and the remaining cells were separately collected for: (1) RNA isolation (gene expression analysis); (2) DNA isolation (DNA methylation analysis); (3) proliferation assay and (4) storage for future use. All steps were repeated for a total of 11 weeks. In parallel with the cetuximab treated cells, we generated controls that received the same correspondent volume of PBS (phosphate buffered saline). Cells were plated in several replicates each time at the same initial density. The replicates were then harvested and pooled to provide enough cells for genetic, epigenetic and proliferation assays. To achieve adequate final cell confluence and number of cells for the experimental analysis of each generation, cetuximab and PBS treated cells were plated in different flask sizes. Cells treated with cetuximab were plated in multiple T75 (75cm^2^) flasks (60,000 cells/flask) that were combined on the eighth day. PBS treated cells were plated in a single T175 (175cm^2^) flask (60,000 cells). This design was selected considering the growth inhibition of the earliest cetuximab generations and to control confluence of the PBS controls at the collection time (**Supplemental Fig. 1**).

### Cell proliferation and colony formation assays

Cell proliferation events were measured using the Click-iT Plus EdU Flow Cytometry Assay Kit Alexa Fluor 488 Picolyl Azide (Life Technologies, Carlsbad, CA) according to manufacturer’s instructions. The cetuximab generations were considered resistant when the frequency of proliferating cells was higher than in the PBS control generations. Proliferation curves were generated using locally weighted polynomial regression (lowess) in R.

Anchorage-independent growth assay was used to further confirm the development of resistance. The parental SCC25 and the late G10 resistant cells were treated with different concentrations of cetuximab 10nM, 100nM and 1000nM. Number of colonies was compared to the same cells treated with PBS. Colony formation assay in Matrigel (BD Biosciences, Franklin Lakes, NJ) was performed as described previously [30].

### Stable SCC25 cetuximab resistant single clones (CTXR clones)

Resistance to cetuximab was induced in an independent passage of SCC25 cells. After resistance was confirmed, single cells were isolated and grown separately to generate the isogenic resistant single cell clones (CTXR). In total, 11 CTXR clones were maintained in culture without addition of cetuximab. With the exception of one clone (CTXR6), all CTXR clones presented substantial survival advantage compared to the parental SCC25, as reported by Cheng et al. (2015) [31]. Each of these clones was authenticated using STR analysis kit GenePrint 10 (Promega) through the JHU-GRCF, as previously published [31].

Proliferation assay was performed to confirm cetuximab resistance in the CTXR clones compared to the parental SCC25. A total of 1000 cells were seeded in 96-well plates in quadruplicate for each condition. PBS or cetuximab (10nM, 100nM or 1000nM) was added after 24 and 72 hours and cells were maintained in culture for 7 days. AlamarBlue reagent (Invitrogen, Carlsbad, CA) at a 10% final concentration was incubated for 2 hours and fluorescence was measured according to the manufacturer’s recommendations (545nm excitation, 590nm emission). Resistance in the CTXR clones was confirmed when the proliferation rates were higher than in the PBS treated SCC25 cells.

### RNA-sequencing (RNA-seq) and data normalization

RNA isolation and sequencing were performed for the parental SCC25 cells (G0) and each of the cetuximab and PBS generations (G1 to G11) and the CTXR clones at the Johns Hopkins Medical Institutions (JHMI) Deep Sequencing & Microarray Core Facility. RNA-seq was also performed for two additional technical replicates of parental SCC25 cell line to distinguish technical variability in the cell line from acquired resistance mechanisms. Total RNA was isolated from a total of 1x10^6^ cells using the AllPrep DNA/RNA Mini Kit (Qiagen, Hilden, Germany) following manufacturer’s instructions. The RNA concentration was determined by the spectrophotometer Nanodrop (Thermo Fisher Scientific, Waltham, MA) and quality was assessed using the 2100 Bioanalyzer (Agilent, Santa Clara, CA) system. An RNA Integrity Number (RIN) of 7.0 was considered as the minimum to be used in the subsequent steps for RNA-seq. Library preparation was performed using the TrueSeq Stranded Total RNAseq Poly A1 Gold Kit (Illumina, San Diego, CA), according to manufacturer’s recommendations, followed by mRNA enrichment using poly(A) enrichment for ribosomal RNA (rRNA) removal. Sequencing was performed using the HiSeq platform (Illumina) for 2X100bp sequencing. Reads were aligned to hg19 with MapSplice [32] and gene expression counts were quantified with RSEM [33]. Gene counts were upper-quartile normalized and log transformed for analysis following the RSEM v2 pipeline used to normalize TCGA RNA-seq data [34]. All RNA-seq data from this study is available from GEO (GSE98812) as part of SuperSeries GSE98815.

### DNA methylation hybridization array and normalization

Genome-wide DNA methylation analysis was performed on the same samples as RNA-seq using the Infinium HumanMethylation450 BeadChip platform (Illumina) at the JHMI Sidney Kimmel Cancer Center Microarray Core Facility. Briefly, DNA quality was assessed using the PicoGreen DNA Kit (Life Technologies) and 400ng of genomic DNA was bisulfite converted using the EZ DNA Methylation Kit (Zymo Research, Irvine, CA) following manufacturer’s recommendations. A total volume of 4μL of bisulfite-converted DNA was denatured, neutralized, amplified and fragmented according to the manufacturer’s instructions. Finally, 12 μL of each sample were hybridized to the array chip followed by primer-extension and staining steps. Chips were image-processed in the Illumina iScan system. Data from the resulting iDat files were normalized with funnorm implemented in the R/Bioconductor package minfi (version 1.16.1) [35]. Methylation status of each CpG site was computed from the signal intensity in the methylated probe (M) and unmethylated probe (U) as a *β* value as follows:

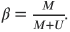

Annotations of the 450K probes to the human genome (hg19) were obtained from the R/Bioconductor package FDb.InfiniumMethylation.hg19 (version 2.2.0). Probes on sex chromosomes or annotated to SNPs were filtered from analysis. The CpG island probe located closest to the transcription start site was selected for each gene. Genes with CpG island probes less than 200bp from the transcription start site were retained to limit analysis to CpG island promoter probes for each gene. Probes were said to be unmethylated for *β* < 0.1 and methylated for *β* > 0.3 based upon thresholds defined in TCGA analyses [34]. All DNA methylation data from this study is available from GEO (GSE98813) as part of SuperSeries GSE98815.

### Hierarchical clustering and CoGAPS analysis

Unless otherwise specified, all genomics analyses were performed in R and code for these analyses is available from https://sourceforge.net/projects/scc25timecourse.

The following filtering criterion for genes from the profiling of the time course data from generations of cetuximab treated cells was used. Genes from RNA-seq data were selected if they had log fold change greater than 1 between any two time points of the same condition and less than 2 between the replicate control samples at time zero (5,940 genes). CpG island promoter probes for each gene were retained if the gene switched from unmethylated *(β* < 0.1) to methylated *(β* > 0.3) in any two samples of the time course (1,087 genes). We used the union of the sets of genes retained from these filtering criteria on either data platform for analysis, leaving a total of 6,445 genes in RNA-seq and 4,703 in DNA methylation.

Hierarchical clustering analysis was performed with Pearson correlation dissimilarities between genes and samples on all retained genes. CoGAPS analysis was performed on both log transformed RNA-seq data and DNA methylation *β* values, independently using the R/Bioconductor package CoGAPS [23] (version 2.9.2).

CoGAPS decomposes a matrix of data **D** according to the model

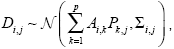

where 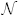 represents a univariate normal distribution, matrices **A** and **P** are learned from the data for a specified number of dimensions *p*, Σ_*i,j*_ is an estimate of the standard deviation of each row and column of the data matrix **D**, and i represents each gene and j each sample. In this decomposition, each row of the pattern matrix **P** quantifies the relative association of each sample with a continuous vector of relative gene expression changes in the corresponding column of **A**. That is each row of P provides a relative magnitudes across samples are called patterns and quantify the separation of distinct experimental conditions. These relative gene weights in the columns of **A** represent the degree to which each gene is associated with an inferred pattern, and called meta-pathways. Together, these matrices provide a low-dimensional representation that reconstructs the signal of the input genomics data. A single gene may have non-zero magnitude in several distinct gene sets, representing the fact that a single gene can have distinct roles in different biological processes (such as immediate therapeutic response and acquired resistance). A recently developed PatternMarker statistics [26] selects the genes that are unique to each of the inferred patterns, and therefore represent biomarkers unique to the corresponding biological process.

In the CoGAPS analysis of the data in this study, the standard deviation of the expression data was 10% of the signal with a minimum of 0.5. The standard deviation of DNA methylation data under the assumption that β values follow a beta distribution is

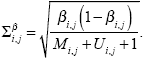

CoGAPS was run for a range of 2 to 10 dimensions *p* for expression and 2 to 5 for DNA methylation. Robustness analysis with ClutrFree [36] determined that the optimal number of dimensions *p* for expression was 5. DNA methylation was run in 4 parallel sets using GWCoGAPS [26]. In DNA methylation, the maximum number of patterns that modeled resistance mechanisms over and above technical variation in replicate samples of SCC25 was three. Gene sets representative of the meta-pathway were derived for each pattern using the PatternMarkers statistics [26]. Comparisons between DNA methylation and gene expression values for PatternMarkerGenes or from CoGAPS patterns and amplitudes were computed with Pearson correlation.

### Gene set analysis of cetuximab resistance signatures, the EGFR network, and pathways

Gene set activity was estimated with the gene set statistic implemented in calcCoGAPSStat of the CoGAPS R/Bioconductor package [23]. Analyses were performed on three gene sets: resistance signatures, gene targets of transcription factors in the EGFR network, and canonical pathways. Resistance signatures were defined based on previous literature. Specifically, in a previous study, CoGAPS learned a meta-pathway from gene expression data corresponding to overexpression of the HRAS^Val12D^ in the HaCaT model of HPV-HNSCC premalignancy. That study associated the CoGAPS HaCaT-HRAS meta-pathway with gene expression changes in acquired cetuximab resistance in the HNSCC cell line UMSCC1 [23]. In the current study, we applied the PatternMarkers statistics [26] to the previously published CoGAPS analysis of these data to derive a gene set from this meta-pathway called HACAT_HRAS_CETUXIMAB_RESISTANCE or HACAT_RESISTANCE. In addition, we searched MSigDB [37] (version 5.2) for all gene sets associated with resistance to EGFR inhibition. In this search, we found the gene sets

COLDREN_GEFITINIB_RESISTANCE_DN and COLDREN_GEFITINIB_RESISTANCE_UP representing resistance to the EGFR inhibitor gefitinib in non-small-cell lung cancer (NSCLC) cell lines [38]. Gene sets of transcription factor targets were obtained from experimentally validated targets annotated in the TRANSFAC [39] professional database (version 2014.1). Canonical pathways were obtained from the C2 set of MSigDB [37] (version 6.1).

### Sources and analysis of additional *in vitro* and human tumor genomics data

Genomics analyses of TCGA were performed on level 3 RNA-seq and DNA methylation data from the 243 HPV-negative HNSCC samples from the freeze set for publication [36]. DNA methylation data was analyzed for the same CpG island promoter probes obtained in the cell line studies. Pearson correlation coefficients were computed in R to associate different molecular profiles.

Additional analysis was performed on Affymetrix Human Genome U133 plus 2.0 GeneChip arrays for the SCC1/1CC8 isogenic cetuximab sensitive and resistant cell line pair described previously (GEO GSE21483 [30]). Additional gene expression data from SCC25 generated from the same platform in the same lab was also used for analysis, using fRMA for normalization [40] to control for batch effects as described previously[41]. Analysis was also performed on gene expression data measured with Illumina HumanHT-12 WG-DASL V4.0 R2 expression beadchip arrays on samples from patients treated with cetuximab from Bossi et al [42], using expression normalization and progression-free survival groups as described in the study. Data were obtained from the GEO GSE65021 series matrix file.

DNA samples from eight human tumor surgical specimen post cetuximab treatment from the sample cohort in Schmitz et al (2015) [43] were obtained for methylation profiling. Specifically, for each tumor one FFPE slide was stained with hematoxylin and eosin and tumor burden was evaluated. When the tumor content was lower than 50%, the adjacent unstained FFPE slides were macrodissected in order to enrich the tumor burden. A double code was assigned to each sample by Biorepository. DNA was then extracted from two unstained slides using the QIAamp DNA FFPE Tissue kit. Briefly, slides were dipped into a xylene bath until paraffin was melted. Then slides were washed with ethanol 100%, and tissue was harvested for extraction with QIAGEN affinity columns. The extracted DNA was quantified by NanoDrop spectrophotometer. DNA methylation was measured with the Illumina MethylationEPIC BeadChip (850K) array. Array data were normalized with NOOB method [44] and converted to virtual 450K arrays using the R/Bioconductor package minifi version 1.22.1 and are available from GEO (in process). Two samples had DNA content below 250ng and clustered separately from the remaining samples. These samples were filtered as low quality and excluded from the analysis, leaving 6 total tumor samples with DNA methylation data. Probes selected for the *in vitro* Illumina 450K DNA methylation data were used for subsequent analyses. Gene expression data from biopsy samples prior to cetuximab treatment and surgical samples post cetuximab treatment were obtained from the previous Schmitz et al [43] study and normalized as described previously [41] and available from GEO (in process).

We performed t-tests and projections in R on the probe that had the highest standard deviation of expression values for each gene. CoGAPS signatures were also projected into these gene expression data using the methods described in Fertig et al. [23] with the ProjectR package version 0.99.15 available from Github (https://github.com/genesofeve/projectR). We also performed t-tests in R to compare the long and short term progression free survival groups based on the values obtained from this projection.

HNSCC samples and patient information collection were approved by the Independent Ethics Committee and the Belgian Health Authorities and conducted in accordance with the Declaration of Helsinki (October 2000).

It was prospectively planned to perform translational research and patients gave their informed consent for repeated biopsies.

## RESULTS

### Prolonged exposure to cetuximab induces resistance

Cetuximab resistance was induced by treating the SCC25 cells for a period of eleven weeks (CTX-G1 to - G11). SCC25 cells treated with PBS were used as time-matched controls (PBS-G1 to -G11). Response to cetuximab was determined by comparing the proliferation rates between CTX and PBS generations. Proliferation of the PBS generations is stable throughout the eleven weeks (G1 to G11). Conversely, proliferation of the CTX generations progressively increases over each week (Figure 1). Relative to the untreated controls, the growth of the treated cells is initially (CTX-G1) inhibited until CTX-G3. Starting at CTX-G4, the absolute proliferation values are equal at this week, but the fit to the data across all time points suggests that the cells become resistant to the anti-proliferative effects of cetuximab and gain stable growth advantages compared to the untreated controls (CTX-G8 to -G11).

**Figure 1.**
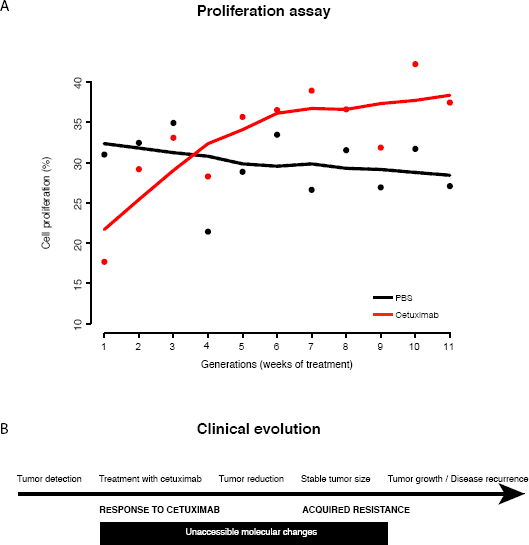
*In vitro* time course reflects clinical evolution of cetuximab response and evolution of acquired resistance. Intrinsic cetuximab sensitive HNSCC cell line SCC25 was treated with cetuximab (red) or PBS (black) for 11 generations to develop acquired resistance. Proliferation assay (flow) of cetuximab treatment (red line) and PBS treated cells (black line) measured cetuximab response for all SCC25 generations (bottom). Treatment response was divided into three stages based upon the measured proliferation rates and clinical stages (top).While proliferation of the PBS generations was stable throughout the eleven weeks, proliferation of the CTX generations progressively increased over each week. Relative to the untreated controls, the growth of the treated cells was initially (CTX-G1) inhibited until CTX-G3. Starting at CTX-G4, the cells became resistant to the anti-proliferative effects of cetuximab and gained stable growth advantages compared to the untreated controls.

Comparison of proliferation rates between generations of CTX treated cells relative to generations of cells treated with PBS enabled us to conclude that cell growth advantages arise from chronic cetuximab treatment and are associated with resistance rather than prolonged cell culturing. We mirrored the changes in proliferation rates with clinical responses seen in HNSCC tumors treated with cetuximab (Figure 1, lower panel). The lower growth rates in CTX-G1 to -G3 may be an equivalent to the initial effects of the clinical treatment when the cancer cells are sensitive to cetuximab and reduction of the tumor size is observable. Even with gain in cell proliferation at CTX-G3 and -G4, our model still corresponds to response to the treatment since the treated generations are not growing more than the controls (clinical stable tumor size). Finally, from CTX-G4 the higher proliferation even with cetuximab treatment is a representation of acquired resistance noticeable in the HNSCC patients as tumor recurrence or increase in tumor size.

Higher proliferation in treated than in untreated cells starts at CTX-G4 and we established this time point to call as the moment at which cetuximab resistance is stably acquired and all subsequent time points continue to develop acquired stable cetuximab resistance. To confirm this hypothesis, we evaluated the ability of the resistant CTX-G10 to anchorage-independent growth. Even under different concentrations of cetuximab, CTX-G10 presents enhanced anchorage-independent growth compared to the parental SCC25 (G0) (twoway Anova with multiple comparisons p-value < 0.01 for each concentration, **Supplemental Fig. 2**), demonstrating the stabilization of cetuximab resistance in later generations.

### Treatment vs. control gene expression changes governs clustering and immediate therapeutic response is confounded with changes from acquired resistance

RNA-seq data for the parental SCC25 cell line (G0) and from each generation of CTX- and PBS-treated cells were collected to characterize the gene expression changes occurring as cells acquired cetuximab resistance. Gene expression changes between treated (cetuximab) and untreated (PBS) cells and over generations of treated cells are apparent in time-ordered RNA-seq data (**Fig. 2A**). Additional clustering analysis of the samples accounting for the treatment time point (generations/columns) (Supplemental Fig. 3) distinguish three clusters of samples: those with cetuximab sensitivity (CTX-G1 to CTX-G3), with early cetuximab resistance (CTX-G4 to CTX-G8), and those with late or stable cetuximab resistance (CTX-G9 to CTX-G11). The group of cetuximab sensitive samples corresponds to the time points at which the CTX generations present lower proliferation rates than the PBS controls (shown in Figure 1). The two groups of samples resistant to cetuximab are represented by a progressive increase in proliferation that is more significant than in the untreated controls (weeks 4 to 8) and by the stabilization in the proliferation rates (weeks 9 to 11), but still higher than in the PBS generations. The expression changes at the distinct time points during development of acquired resistance are shared among numerous genes. Although the clustering was able to separate cetuximab from PBS treated cells, it was not possible to discriminate the alterations related to an immediate therapeutic response (not relevant to the resistant phenotype) from resistance-specific gene expression changes.

**Figure 2.**
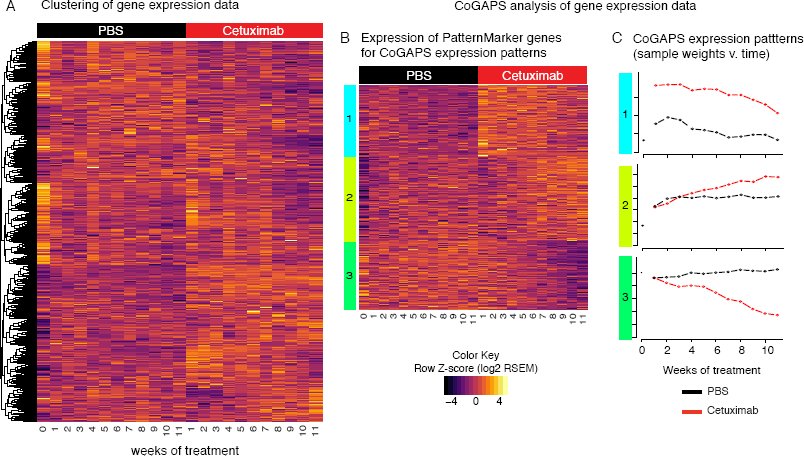
CoGAPS analysis identifies signatures of resistance to EGFR inhibitors and separate resistant and control generations. **(A)** Heatmap of gene expression values in 11 generations of SCC25 cells treated with 100nM of cetuximab (red columns) to acquire resistance and with PBS as control (black columns). **(B)** Heatmap of gene expression values for PatternMarker genes identified with CoGAPS analysis of gene expression data from 11 generations of SCC25 cells treated with PBS as control (black columns) and with 100nM of cetuximab (red columns) to acquire resistance. Rows are colored according to which CoGAPS pattern the PatternMarker statistic assigned each gene, and sorted by the PatternMarker statistic. **(C)** CoGAPS patterns inferred from gene expression data over generations of PBS control (black lines) or treatment with 100nM of cetuximab (red lines).

Similar separation of the phases of cetuximab response is observed in clustering analysis of gene signatures previously described in HNSCC and NSCLC cell line models resistant to cetuximab or gefitinib (anti-EGFR small molecule), respectively [38,41] (**Supplemental Fig. 4**). For these genes, changes during early phases of resistance clusters for CTX-G4 to CTX-G6 as distinct from later generations CTX G7-11. Nevertheless, these signatures also cluster samples with gene expression changes at early phases (CTX-G1 to G3) as distinct from samples from PBS treated generations. However, these analyses were insufficient to quantify the relative dynamics of genes associated with immediate response to therapy or subsequent acquired resistance.

### CoGAPS analysis of gene expression distinguishes patterns of acquired resistance from immediate therapeutic response

To define gene expression signatures for treatment effect and cetuximab resistance, we applied CoGAPS [26] Bayesian matrix factorization algorithm to the time course gene expression data. CoGAPS decomposes the input data into two matrices: a pattern matrix with relative sample weights along rows and an amplitude matrix with relative gene weights along columns. Each row of the pattern matrix quantifies the extent of transcriptional changes within the genes in the corresponding column of the amplitude matrix, and provides a low dimensional representation of the biological process in that data. In this analysis, we identified five CoGAPS patterns (Expression Patterns - EP) (**Supplemental Fig. 5**) in the time course gene expression dataset. A gene can have high amplitude in multiple patterns, modeling multiple regulation of genes but complicating visualization of the inferred patterns. A recently developed PatternMarker statistic defines genes that are uniquely associated with each of these patterns. Limiting the heatmap to these genes enables visualization of the dynamics of gene expression changes in our time course dataset (**Supplemental Fig. 5**).

In this heatmap of CoGAPS PatternMarker genes, we observe that only three patterns (EP1, EP2 and EP3) distinguish the experimental conditions (cetuximab vs. PBS) (top three patterns on Supplemental Fig.5). The other two patterns, EP4 and EP5, represent changes in gene expression from the parental cell lines and subsequent generations or an expression pattern that is constant and corresponds to signature of highly expressed genes (lower two patterns on Supplemental Fig. 5), respectively. We note that highly expressed genes associated with EP5 may also have dynamic changes due to treatment, and are filtered in the PatternMarker analysis of all patterns in Supplemental Figure 5. EP4 represents expression changes between treated cells and the parental cell line, which have a technical effect on gene expression. Notably, even the exclusion the flat pattern for highly expressed genes (EP5) still retains highly expressed genes in the signature. Retaining the technical pattern (EP4) in this calculation filters genes with expression changes from technical artifacts in the experimental conditions. The resulting set of PatternMarker genes for EP1 - EP3 enable visualization of the expression changes dynamics that are associated with cetuximab response (**Fig 2B**) and allow the definition of a gene signature associated with that response (**Supplemental Table 1**).

Similar to the separation seen with clustering (**Supplemental Fig. 5**), the CoGAPS pattern EP1 distinguishes cetuximab from PBS at every generation (**Fig. 2B** and **Fig. 2C**, top). These genes present an immediate transcriptional up-regulation in response to cetuximab treatment. Gene set analysis to determine the function of CoGAPS patterns was performed with an enrichment analysis on all gene weights in the amplitude matrix obtained from the CoGAPS analysis. By performing the analysis on gene weights and not only the PatternMarker genes, as shown in **Fig. 2B**, we account for multiple regulation of genes in pathways. Specifically, we performed gene set analysis on published resistance signatures [38,41], transcription factors previously associated with the EGFR signaling network during cetuximab response in HNSCC [39,41], and canonical pathways from MSigDB [37,45] (**Supplemental Fig. 6; Supplemental Table 2**). Gene set analysis confirms that published resistance signatures [38,41] are significantly enriched in EP1 (**Supplemental Fig. 6**; one-sided p-values of 0.002 and 0.003 for resistance gene sets COLDREN_GEFITINIB_RESISTANCE_DN and HACAT_HRAS_CETUXIMAB_RESISTANCE, respectively). However, the transcriptional changes in this pattern are not associated with acquired resistance to cetuximab, and even decrease modestly as resistance developed. Further, enrichment by transcription factor AP-2alpha targets *(TFAP2A;* one-sided p-value of 0.05) confirms previous work indicating that transcription by AP-2alpha is induced as an early feedback response to EGFR inhibition [39]. There are 84 significant canonical pathways from MSigDB, including notably pathways associated with the immune system, extracellular matrix, ERBB4 signaling, and VEGF signaling (**Supplemental Table 2**). Based upon these findings, we concluded that EP1 is associated with immediate response to cetuximab although it includes genes that are also associated with cetuximab resistance in previous studies.

The second CoGAPS expression pattern (EP2) quantifies divergence of the cetuximab treated cells from controls at generation CTX-G4 (**Fig. 2B** and **Fig. 2C**, middle) which is the time point that cetuximab treated cells present significant and stable growth advantage over PBS controls (**Fig. 1**). Therefore, EP2 contains gene expression signatures associated consistently with the development of cetuximab resistance. Gene set statistics of transcription factor targets of EGFR on CoGAPS gene weights are significantly down-regulated in this acquired resistance pattern (**Supplemental Fig. 6**). One striking exception is c-Myc, which trends with acquired resistance (p-value of 0.06), consistent with the role of this transcription factor in cellular growth. Resistance signature COLDREN_GEFITINIB_RESISTANCE_DN is significantly down-regulated in EP2 (p-value of 0.04). There are 32 statistically significant canonical pathways associated with this pattern, including notably telomerase, PI3K, and cell cycle pathways (**Supplemental Table 2**).

The third CoGAPS expression pattern (EP3) represents a gradual repression of gene expression with cetuximab treatment (**Fig. 2B** and **Fig. 2C**, bottom). This expression pattern trends to significant enrichment in the COLDREN_GEFITINIB_RESISTANCE_DN resistance signature (**Supplemental Fig. 6**, one-sided p-value 0.12) and down-regulated in the HACAT_HRAS_CETUXIMAB_RESISTANCE resistance signature (Supplemental Fig. 6, one-sided p-value 0.09). This confirms that EP3 is associated with repression of gene expression during acquired cetuximab resistance. There are also 29 statistically significant canonical pathways associated with this pattern, including cell lineage, metabolic, WNT, and GSK3 pathways (**Supplemental Table 2**).

### Changes in DNA methylation inferred with CoGAPS are associated with resistance to cetuximab, but not the immediate response to treatment observed in gene expression

To determine the timing of the methylation changes associated with acquired resistance, we also measured DNA methylation in each cetuximab generation of SCC25 cells and PBS controls (**Fig. 3A**). Application of the CoGAPS matrix factorization algorithm with the PatternMarker statistics to the methylation data reveals a total of 3 methylation patterns (MP) (**Fig. 3BC; Supplemental Table 1**): gradual increase of DNA methylation in controls (MP1, Fig. 3B middle); rapid demethylation in CTX generations starting at CTX-G4 (MP2, Fig. 3B bottom); and rapid increase in DNA methylation in CTX generations starting at CTX-G4 (MP3, Fig. 3B top). In contrast to the gene expression data, there is no immediate shift in DNA methylation resulting from cetuximab treatment. Gene set analysis was performed on canonical pathways from MSigDB (**Supplemental Table 3**), and found 26 statistically significant pathways for MP1, 29 for MP2, and 27 for MP3. In contrast to gene expression, the majority of canonical pathways is shared by the three methylation patterns and include notably the cytokine (PID-IL8-CXCR2 and IL8-CXCR1 pathways) and FGFR (Reactome Signaling by FGFR3 mutants) signaling pathways.

**Figure 3.**
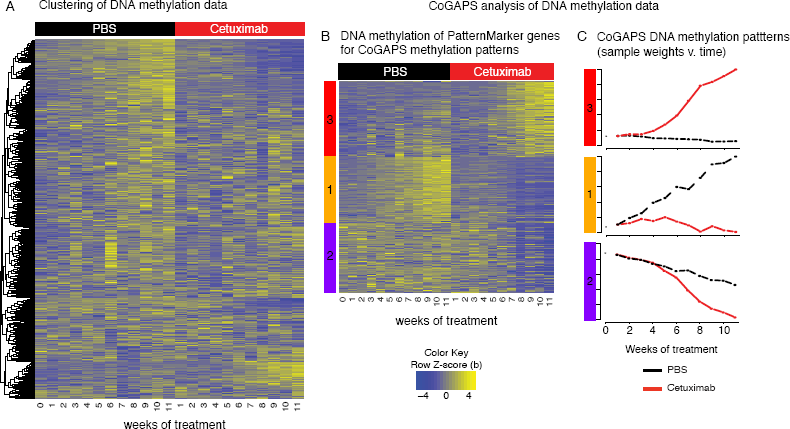
Dynamics of DNA methylation alterations and association with gene expression patterns in acquired cetuximab resistance. **(A)** Heatmap of DNA methylation values in 11 generations of SCC25 cells treated with PBS as control (black columns) and with 100nM of cetuximab (red columns) to acquire resistance. **(B)** Heatmap of DNA methylation values for genes selected by CoGAPS DNA methylation patterns analysis in the same SCC25 cetuximab and PBS generations. **(C)** CoGAPS patterns inferred from DNA methylation data over generations of PBS control (black lines) or treatment with 100nM of cetuximab (red lines).

Comparing the CoGAPS patterns from gene expression and DNA methylation reveals strong anti-correlation between gene expression and DNA methylation in resistant patterns (**Supplemental Fig. 7A**). We observed that the gene expression changes associated with acquired resistance occur gradually and are evident in early generations (**Fig. 2C**). The DNA methylation is consistent in cetuximab treatment and control PBS in DNA methylation patterns MP2 and MP3 during early generations. Then, rapid accumulation in DNA methylation changes starts after generations CTX-G4 and CTX-G5 in both MP2 and MP3 (**fig. 3C**), concurrent with the onset of the observed growth advantage over the PBS control (**Fig. 1**).

While the patterns themselves are anti-correlated, the gene weights that define meta-pathways and are inferred in the amplitude matrix corresponding to each pattern with CoGAPS are not (**Supplemental Fig.7B**). We also observed little overlap between the PatternMarker genes from methylation patterns and gene expression. Changes in DNA methylation are delayed relative to those of gene expression in acquired cetuximab resistance as can be noted in **Fig. 4**, where direct comparison of the expression and methylation patterns previously shown (**Fig. 2C** and **3C**, respectively) enable visualization of the time point when changes between cetuximab and PBS generations are significant in each pattern. These dynamics explain the discrepancy between the genes associated with each pattern and suggest that DNA methylation stabilizes the gene expression signatures crucial to the maintenance of acquired cetuximab resistance.

**Figure 4.**
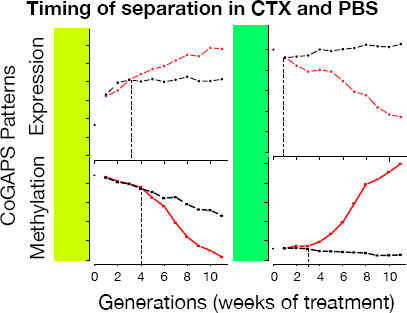
DNA methylation and expression CoGAPS patterns demonstrate delayed onset of epigenetic changes in acquired resistance. CoGAPS patterns for gene expression (top) and DNA methylation (bottom) of patterns associated with acquired cetuximab resistance in SCC25 cetuximab generations (red) relative to PBS generations (black). Vertical dashed line represents time at which patterns for SCC25 generation separated from pattern for PBS generations. The timing of methylation changes distinguishing cetuximab resistant generations was delayed in DNA methylation relative to that of gene expression.

### Gene expression and methylation profile of SCC25 single-cell clones with acquired cetuximab resistance demonstrates cell heterogeneity

The little overlap between the gene expression and DNA methylation PatternMarker genes and non-specific DNA methylation pathways may arise due to the development of different resistant sub-clones with specific gene signatures of acquired resistance in bulk data. In order to address this issue and to delineate whether our presumptive drivers resulted from clonal expansion of resistant cells or from the development of new epigenetic alterations to drive resistance, we measured DNA methylation and gene expression on a panel of eleven isogenic stable cetuximab resistant clones (CTXR1 to CTXR11) derived from SCC25 cells in a previous study [31]. Despite being derived from the parental SCC25 cells after chronic exposure to cetuximab, the CTXR clones and the time course generations display widespread differences. Significantly greater heterogeneity is observed among the CTXR clones in both expression and methylation profiles (**Supplemental Fig. 8** and **9**, respectively) and cellular morphology (**Supplemental Fig. 10**). **Fig. 5A and 5B** demonstrate that higher heterogeneity among single cell clones is also observed in the epigenetically regulated PatternMarker genes from the CoGAPS analysis. These results suggest that different mechanisms of resistance may arise in the same HNSCC cell line as a result of intra-heterogeneity, resulting in the detection of a wide-range of expression signatures with higher or lower correlation with the methylation profile depending on the size of each specific cell population.

**Figure 5.**
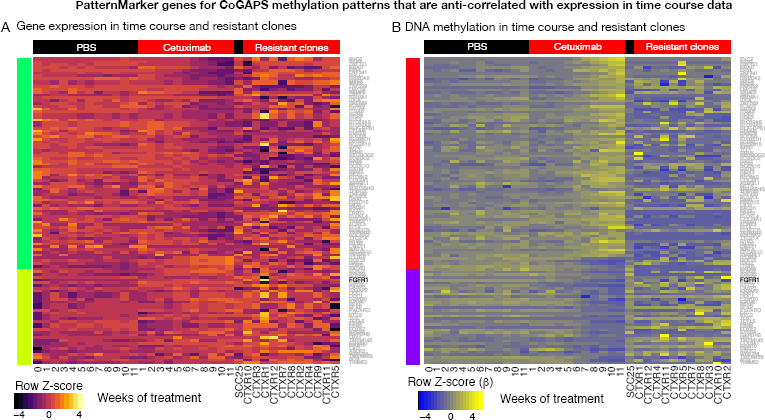
Clonal heterogeneity does not reflect signature of epigenetically regulated genes observed in bulk time course analysis. **(A)** Heatmap of gene expression values for DNA methylation PatternMarker genes for acquired resistance that were anti-correlated between expression and DNA methylation (**Fig. 4**). Data includes 11 generations of SCC25 cells treated with PBS as control (black columns labeled PBS) and with 100nM of cetuximab (red columns labeled cetuximab) to acquire resistance and gene expression data from independent, stable cetuximab resistant clones in absence of cetuximab treatment (CTX resistant clones). Gene expression heatmap on a red-yellow scale indicated in the color key. **(B)** Heatmap of DNA methylation data in conditions described in **(A)**, on a blue-yellow scale indicated in the color key.

### *FGFR1* over-expression and demethylation are associated with acquired cetuximab resistance in the time course and in stable cetuximab resistant clones

To ascertain potential drivers of the stable cetuximab resistant phenotype induced by DNA methylation, we defined genes that are PatternMarkers [26] of the DNA methylation patterns associated with stable acquired cetuximab resistance (MP2 and MP3). We then applied correlation analysis to determine genes that were epigenetically regulated. Specifically, we performed correlation analysis between DNA methylation and gene expression for each of the DNA methylation PatternMarker genes (**Fig. 5**). This analysis identified *FGFR1* as one of the genes with significant anti-correlation between expression and methylation, suggesting potential epigenetic regulation during cetuximab resistance acquisition. This finding is consistent with previous studies that associate differential expression of *FGFR1* with resistance to EGFR inhibitors, including cetuximab, in HNSCC and other tumor types *in vitro* and *in vivo* [27,46-48]. However, none of these studies demonstrate an association between *FGFR1* up-regulation and hypomethylation. Given the tight temporal regulation of these genes and the previous work on *FGFR1,* we hypothesize that this set of genes represents epigenetic drivers of acquired resistance.

We hypothesize that epigenetically regulated genes shared along the time course patterns and resistant single-cell clones might implicate common mechanisms acquired during evolution of the stable resistance phenotype. To test this assumption, we also performed correlation analysis for each of the epigenetically regulated genes in our resistant set (**Fig. 5**) in the resistant single cell clones and parental cell lines. Nine of the epigenetically regulated PatternMarker genes also have significantly anti-correlated gene expression and DNA methylation in the stable cetuximab resistant clones (**Supplemental Fig. 11**). Of these, only *FGFR1* is demethylated and re-expressed in a CTXR clone relative to the parental SCC25 cell line (**Fig. 6**). In this analysis, over-expression and de-methylation of *FGFR1* expression occurs in only one of the resistant clones (CTXR10). This clone is one of the fastest growing under cetuximab treatment (**Supplemental Fig. 12**). This observation suggests that the bulk data from the time course captured clonal outgrowth of a cetuximab resistant clone with similar molecular features *(FGFR1* hypomethylation) to CTXR10, and that clonal outgrowth is the dominant mechanism of resistance in our model.

**Figure 6.**
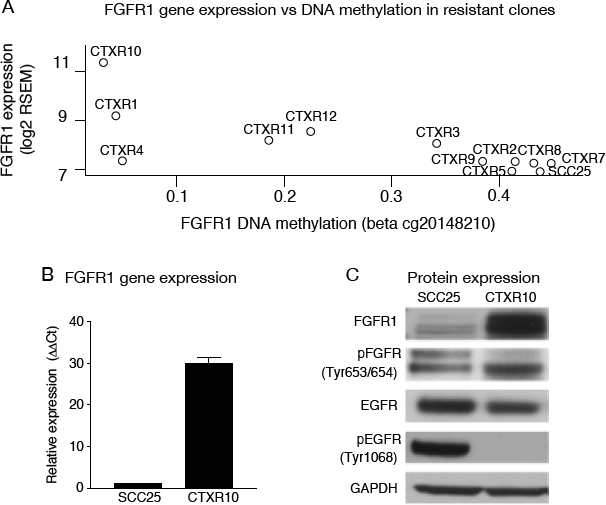
Overexpression and de-methylation of ***FGFR1*** in acquired cetuximab resistance is confirmed in stable SCC25 resistant clones. **(A)** Expression of ***FGFR1*** gene expression relative to DNA methylation in stable cetuximab resistant clones. **(B)** QRT-PCR of ***FGFR1*** gene expression in CTXR10 relative to the parental cell line (greater than 30 fold change). **(C)** Western blot comparing FGFR1, phosphor-FGFR1, EGFR, and phospho-EGFR in CTXR10 relative to the parental SCC25 cell line. In the resistant cell clone, increased levels of FGFR1 were associated with increased levels of phospho-FGFR1 and decrease in EGFR and phospho-EGFR.

### Observed *FGFR1* dynamics *in vitro* recapitulates relationships from *in vivo* tumor genomics and acquired cetuximab resistance

In order to confirm that the mechanisms we found with our *in vitro* approach are present in HNSCC samples pre- and post-cetuximab treatment, we further investigate the pattern of expression and methylation of *FGFR1* and *EGFR* in publicly available datasets. Using gene expression and DNA methylation data from The Cancer Genome Atlas (TCGA) for 243 HPV-negative HNSCC pretreatment samples [36], we verified that the up-regulation of *EGFR* and *FGFR1* is not concomitant (Pearson correlation coefficient = −0.06, p value = 0.33, **Fig. 7A**). Additionally, the negative correlation of *FGFR1* gene expression and DNA methylation status is statistically significant (Pearson correlation r of −0.32, p value < 0.0001, **Fig. 7B**), suggesting that *FGFR1*transcription is associated with demethylation in some HPV-negative HNSCC tumors. Since there is no treatment information available for the TCGA dataset, we could not make assumptions related to cetuximab resistance and whether *FGFR1* methylation is a consequence of the treatment. To assess this question, we collected new DNA methylation data for six HNSCC tumors after cetuximab treatment from a cohort of HNSCC tumor samples described previously [43]. All six samples have low DNA methylation values for *FGFR1* (β-values ranging from 0.04 to 0.08, with a mean of 0.05), suggesting that the gene is unmethylated in these samples. While there was insufficient DNA to quantify DNA methylation prior to treatment in these patients, *FGFR1* gene expression increases after treatment in four of the six tumor samples (**Supplemental Figure 13**). While this cohort is small, these data and TCGA suggest that *FGFR1* methylation is potentially associated with its reexpression in HNSCC tumor samples in response to cetuximab treatment.

**Figure 7.**
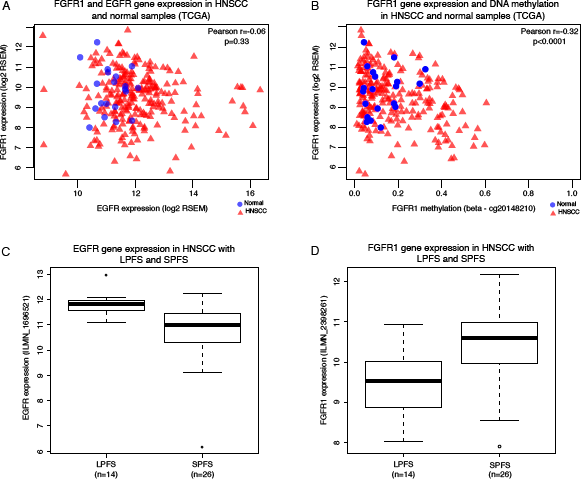
FGFR1 gene expression and DNA methylation patterns were confirmed in independent HNSCC tumor samples datasets. **(A)** Scatter plot of gene expression for ***EGFR*** and ***FGFR1*** in HPV-negative HNSCC samples from TCGA demonstrated that only a few HNSCC cases present increased levels of both genes and that there is no significant correlation between the expression of both genes concomitantly. **(B)** DNA methylation of ***FGFR1*** was anti-correlated with ***FGFR1*** expression in HPV-negative HNSCC TCGA samples, suggesting that up-regulation of FGFR1 might be a result of promoter hypomethylation in primary tumors. **(C) *EGFR*** expression was significantly overexpressed in a group of HNSCC patients with long progression free survival relative to patients with short progression free survival in gene expression data from Bossi et al. **(D) *FGFR1****38* was significantly overexpressed in patients with short progression free survival relative to patients with long progression free survival in this same dataset.

To determine whether *FGFR1* is associated with cetuximab response, we used gene expression data from HNSCC patients before cetuximab treatment available from Bossi et al. [42]. After follow-up, the patients were separated in short- (SPFS, median 3 months survival) or long-progression-free survival (LPFS, median 19 months survival) according to time of recurrence or metastasis development. Using this dataset, we confirmed that *EGFR* expression in SPFS is significantly lower than in the LPSF group (**Fig. 7C**) (log fold change −1.0, t-test p-value 0.0003). The opposite was observed for *FGFR1,* with overexpression in SPFS vs. LPSF (**Fig. 7D**, log fold change 0.9, t-test p-value 0.003). However, Bossi et al. [42] study lacks DNA methylation data to assess whether *FGFR1* was epigenetically regulated in these samples. Most patients with SPFS in this dataset also had intrinsic resistance to cetuximab, instead of acquired resistance studied in our *in vitro* model. Nonetheless, these findings suggest that similar molecular mechanisms may contribute to both mechanisms (intrinsic and acquired) of cetuximab resistance in HNSCC.

### CoGAPS signatures of resistance and therapeutic response replicate in an independent *in vitro*system and significantly stratified patient samples with long vs. short progression free survival

To further illustrate that the results are reflective of HNSCC in a general fashion, we evaluated the behavior of the two additional cell lines and human tumors in the CoGAPS signatures using gene expression data available from previously published studies. The HNSCC cell lines SCC1 and 1CC8 were chosen as the cetuximab resistant 1CC8 was generated from the cetuximab sensitive SCC1 in a similar protocol used to establish the single cell clones [30]. Data from SCC25 was also included as a reference. It is important to note that the treatment time for the SCC1 and 1CC8 pair is on the order of hours vs. weeks, as used to generate the time course data. By projecting these data into the CoGAPS signatures, the relationship between the sensitive SCC1 and resistant 1CC8 recapitulates the relationship between PBS and CTX time course generations (respectively) in treatment driven signatures (**Fig. 8A,B,E,F**). Conversely, CoGAPS signatures related to culture specific conditions failed to produce meaningful differences between the lines (**Fig. 8C,D**). Projections of the expression patterns from CoGAPS into the cell line gene expression data were also anti-correlated with projections of the methylation signatures in these same data. Gene expression data from HNSCC tumors from patients prior to their treatment with cetuximab described in Bossi et al [42] were also analyzed. Projection into both the CoGAPS signatures of resistance and therapeutic response significantly stratified long vs. short progression free survival (p-value=5.2x10^−5^ and 3.1x10^−3^, respectively, (**Fig. 8G,H**). Conversely, projection in to the CoGAPS signature associated with culturing was not significant (p-value = 0.50, Figure 8I).

**Figure 8.**
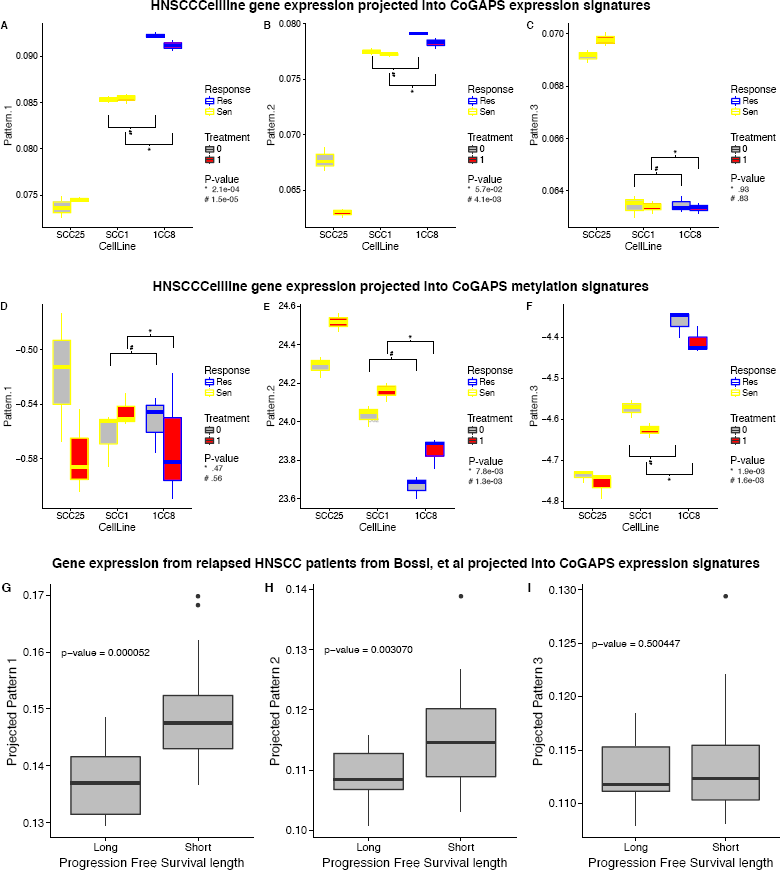
CoGAPS gene signatures confirmed in independent in vitro and in vivo expression data. **(A-C)** Box plots of sample weights for HNSCC cetuximab sensitive cell lines SCC25, SCC1 and resistant 1CC8 in CoGAPS expression signatures. **(D-F)** Box plots of sample weights for HNSCC cetuximab sensitive cell lines SCC25, SCC1 and resistant 1CC8 in CoGAPS methylation signatures. **(G-H)** Box plots of tumor gene expression profiles from relapsed HNSCC patients projected into CoGAPS expression signatures. Patient samples are stratified by length of progression free survival.

## DISCUSSION

Although numerous short time course genomics studies of therapeutic response have been performed [49–51], this is the first time that genetic and epigenetic changes were measured for a prolonged exposure (11 weeks) to a targeted therapeutic agent. Using our novel robust time course integrated analysis approach, we characterized the molecular alterations during the development of acquired cetuximab resistance using a HNSCC *in vitro* model. Cell proliferation, gene expression, and DNA methylation high throughput analysis were performed weekly in equivalent cultures (cetuximab and PBS control generations) as resistance developed. Over the course of 11 weeks, it was possible to compare treated (CTX) and untreated (PBS) cells grown under the same conditions. Applying robust bioinformatics algorithms we discriminated changes associated with acquired resistance from those related to adaptive response to the cell culturing process and treatment. The SCC25 cell line model was the chosen since this is one of the only two HNSCC cell lines previously used to generate isogenic cetuximab resistant cell lines [10]. However, this is the first study to our knowledge to enable characterization of the transcriptional and epigenetic dynamics at the early phases of therapeutic resistance, which cannot be measured in patients due to the complexity of early detection of resistance and obtaining repeated biopsy samples.

Determining the dynamics of the molecular alterations responsible for resistance requires integrated, time course bioinformatics analysis to quantify these alterations. Based upon previous performance of Bayesian, non-negative matrix factorization algorithms in inferring dynamic regulatory networks for targeted therapeutics [49,50], we selected CoGAPS [23] for analysis of gene expression data from our time course experiment. CoGAPS have already proven highly effective in relating gene expression changes to patterns related to EGFR inhibition [41], perturbation of nodes in the EGFR network [39], and time course dynamics of targeted therapeutics. In this dataset, CoGAPS analysis of gene expression data from cetuximab resistant clones distinguished the patterns for immediate gene expression changes and patterns for long-term changes associated with acquired resistance. Gene expression signatures for resistance to EGFR inhibitors in two additional cell lines (one HNSCC and one NSCLC) from previous studies [38,41] were significantly enriched in both types of CoGAPS patterns. Since these previous resistance signatures were learned from case-control studies without multiple time point measurements, we concluded that our time course data is instrumental in discriminating the signatures of immediate therapeutic response from signatures of acquired resistance.

In spite of the complexities of the data integration, the weight of each sample in patterns inferred by CoGAPS reflects the dynamics of the process in each data modality. These patterns are learned completely unsupervised from the data, and do not require any gene selection or comparison between time points relative to any reference control. The CoGAPS analysis of the time course data demonstrates that applying matrix factorization algorithms for genomics can reconstruct signals associated with phenotypes from time course, omics data. The genes associated with CoGAPS patterns had weights that were non-zero in multiple patterns. The PatternMarker statistics [26] enables further selection of the genes that are uniquely associated with each pattern. Creating a heatmap of the genomics profiles for these genes enabled novel, heatmap-based visualization of the temporal dynamics in the omics data. CoGAPS analysis of gene expression data contains a flat pattern (EP5), which includes all highly expressed genes. These genes may also change in association with the experimental conditions, albeit to a lesser degree than lowly expressed genes. Because the PatternMarker statistics includes genes that are uniquely associated with each inferred biological process, these highly expressed genes would be filtered from associations with the dynamic conditions. To include these genes in the signatures defined in this study, EP5 was filtered from the calculation of the PatternMarker statistics. Such filtering process is not required for heatmap-based visualization and filtering of flat patterns is recommend when defining gene signatures containing sets of genes that are most strongly associated with dynamic changes. Patterns that reflect technical artifacts in the data, such as EP4, should be retained in the PatternMarker analysis to limit the signatures associated with inferred processes to retain only biologically relevant genes. We note that these PatternMarker statistics are similar to the D-scores proposed in Zhu et al [52] and that application of this statistic may require similar filtering to retain highly expressed genes. In the case of DNA methylation, these PatternMarker genes also include genes representing driver alterations in resistance. The DNA methylation data did not require filtering when applying the PatternMarker statistics since no flat pattern was detected. However, transcriptional regulation by epigenetic alterations or in pathways involves simultaneous co-regulation of multiple genes. This co-regulation is reflected in the reuse of genes in CoGAPS gene weights associated with each pattern. Therefore, estimates of pathway dynamics from transcriptional data require accounting for all genes with gene set enrichment statistics instead of the PatternMarker statistics. Thus, we hypothesize that the PatternMarker statistics is robust for visualization and biomarker identification. On the other hand, gene set enrichment of the CoGAPS gene weights corresponding to each pattern and stored in the amplitude matrix are essential for characterization of functional alterations in pathways.

Collecting treated and untreated cells to obtain paired measurements of methylation and gene expression enabled us to evaluate whether changes in DNA methylation impact gene expression. Including a PBS control at every time point also enabled the discrimination of the changes that result from an adaptive response to therapy from changes that result from maintaining cells in culture. CoGAPS analysis of DNA methylation data denotes only changes associated with acquired resistance, in contrast to the immediate expression changes observed with cetuximab treatment. Thus, while therapeutic response can drive massive changes in gene expression, only the subset of expression changes associated with the development of resistance have corresponding epigenetic signatures, suggesting that the methylation landscape is important for the development of acquired resistance. The CoGAPS patterns in gene expression that are associated with acquired cetuximab resistance gradually change over the time course (EP1 and EP2). On the other hand, the CoGAPS patterns for DNA methylation changes have a sharp transition at the generation at which resistance is acquired (CTX-G4). These patterns (MP1 and MP2) reflect a delayed but more rapid change in DNA methylation. The time delays between alterations in DNA methylation and gene expression pose a further computational challenge for integrated, time course genomics analyses. The vast majority of integrated analysis algorithms assume one-to-one mapping of genes in different data platforms or seek common patterns or latent variables across them [53]. Such approaches would fail to capture the early changes from cetuximab treatment that impact only gene expression; time delays between DNA methylation and gene expression patterns; and different gene usage in each pattern. It is essential to develop new integrated algorithms to simultaneously distinguish both patterns that are shared across data types and that are unique to each platform. For time course data analysis, these algorithms must also model regulatory relationships that may give rise to timing delays, such as epigenetic silencing of gene expression. However, as we observed with the unanticipated changes in DNA methylation following and not preceding gene expression, they must also consider delays resulting from larger phenotypic changes such as the stability of the therapeutic resistant phenotype.

The relative timing of change in DNA methylation and gene expression is consistent with previous observations that gene expression changes precede DNA methylation alterations in genes critical for cancer progression. *P16^INK4A^* and *GSTP1* are tumor suppressor genes for which transcription silencing was found to occur prior to DNA hypermethylation and chromatin changes. The temporal delay observed between expression and methylation patterns in our time course provides transcriptome wide evidence of this phenomenon. Specifically, that epigenetic changes are necessary to stabilize gene expression aberrant profile and will be followed by modification into a silenced methylation state, resulting in tumor progression [54,55]. Our integrated RNA-seq and DNA methylation analysis corroborates the fact that gene expression changes occur earlier to epigenetic alterations and suggests that DNA methylation is essential to maintain the changes in gene expression in this acquired cetuximab resistance model. Additional time course data tracing other *in vitro* and *in vivo* models of HNSCC are essential to generalize the relative timing of molecular changes, and thus mechanisms of gene regulation, associated with acquired therapeutic resistance. Future investigation into the chromatin remodeling mechanisms will also test whether chromatin alterations follow the changes in expression and occur in combination with altered methylation patterns to drive epigenetic regulation of resistance.

Besides the immediate changes in gene expression followed by the gradual methylation switch, it is also interesting to note these effects in the proliferation rates during the 11 weeks of treatment. Initially, the proliferation of the population of cetuximab sensitive cells is slower when compared to the untreated controls, reflecting therapy effectiveness. Early and progressively, the cells develop molecular changes to overcome the EGFR blockade. However, this process starts in just a small number of clones and the increase in proliferation is still not enough to surpass the growth rate of the untreated cells. As soon as the population of resistant cells is larger than the number of sensitive cells, the proliferation rate is now higher than in the untreated controls. At some point, we observe the stabilization of the proliferation rates in the cetuximab treated cells probably due to the fact the culture is now dominated by the population of resistant clones. Although stable, proliferation rate of these resistant clones is significantly higher than that of the untreated cells. This increased proliferation rate is consistent with the rapid increase in tumor volume observed clinically once patients develop resistance to therapy. Tracing the increase in the population of resistant cells and their proliferation rates *in vivo* require novel techniques to biopsy or image tumors at intermediate time points of treatment.

In a recent study, gene expression changes were associated with a transient resistant phenotype present in melanoma cell lines prior to vemurafenib administration [56]. Once the melanoma cells were exposed to the drug, additional changes in gene expression are detected and are later accompanied by changes in chromatin structure [56]. These findings, together with our time course observations, suggest that in the heterogeneous tumor environment the existence of some cells expressing specific marker genes can trigger cellular reprogramming as soon as the targeted therapy is initiated. Upon drug administration, the number of genes with aberrant expression increases, and is followed by other epigenetic and genetic changes that will shift the transient resistant state into a stable phenotype. This finding on acquired resistance development could dramatically change the course of treatment with targeted therapeutic agents. The precise characterization of resistant gene signatures and their timing are crucial to determine the correct point during the patients’ clinical evolution to introduce alternative therapeutic strategies. This way, secondary interventions would start before the stable resistant phenotype is spread among the tumor cells resulting in prolonged disease control and substantial increase in overall survival.

Among the genes we identified with the canonical relationship between expression and methylation, *FGFR1*present with increased gene expression accompanied by loss of CpG methylation. *FGFR1* is a receptor tyrosine kinase that regulates downstream pathways, such as PI3K/AKT, and RAS/MAPK, that are also regulated by *EGFR* [57]. Its overexpression has been previously associated with resistance to EGFR inhibitors in other cancer types including HNSCC [27–29]. To our knowledge this is the first study showing that epigenetic alterations are associated with changes in *FGFR1* expression in HNSCC during the development acquired cetuximab resistance. *FGFR1* up-regulation combined with promoter hypomethylation was previously described in rhabdomyosarcomas [58]. Other studies described that *FGFR1* increased levels is a common feature in different tumor types, such as glioblastoma [59] and cancers of the breast [60], lung [61], prostate [62], bladder [63], ovarian [64], colorectal [27] and HNSCC [29,65,66]. *FGFR1* is involved in resistance mechanisms against EGFR inhibitors [27,46-48], such as cetuximab and gefitinib. Together, the TCGA and Bossi et al. datasets analysis corroborate our findings that *FGFR1* gene expression is regulated by epigenetic changes in HNSCC. Altogether, the epigenetic alteration of *FGFR1* represents a candidate biomarker of resistance to cetuximab and further studies are critical to identify combination therapies for HNSCC patients that develop acquired cetuximab resistance.

The increased levels of FGFRs and FGFs are believed to play a role in an autocrine mechanism in HNSCC and NSCLC cell lines with intrinsic resistance to the EGFR inhibitor, gefitinib. Using public available gene expression microarray datasets, Marshall et al. [47] and Marek et al. [46] verified concomitant increased levels of FGFRs and their specific FGFs ligands. Particularly, FGFR1 and FGF2 up-regulation were observed in the same resistant cell lines and hypothesized to be the mechanism behind resistance. This was corroborated by functional experiments showing that cells treated with pan-FGFR inhibitor were less prone to anchorage-independent growth. Also *FGF2* silencing or FGFR1 inhibition resulted in phospho-ERK decreased expression that was restored when FGF2 was added to the culture, suggesting that an autocrine

FGF-FGFR pathway is one of the mechanisms of resistance to gefitinib. However, the cell lines evaluated in both studies were intrinsically resistant to gefitinib. In our model, we induced resistance to cetuximab and observed *FGFR1* gain of expression and significant anti-correlation with the DNA methylation. We additionally evaluated the expression of other FGFRs and FGFs that were identified by the PatternMarker statistics. Although it is not found as a PatternMarker in our analysis pipeline, *FGF2* is up-regulated in the cetuximab generations when compared to the PBS generations as observed with *FGFR1* (**Supplemental Fig.14**). Thus, our data corroborates and extends this previous evidence from intrinsically resistant lines that one of the mechanisms driving resistance to EGFR inhibition is the FGF-FGFR autocrine pathway. This observation adds another evidence that the computational approach used in this study is robust once it is capable of identifying mechanisms previously described in other models resistant to EGFR blockade.

Our previously developed bioinformatics algorithms for the identification of gene expression and epigenetic patterns progression over time proved to be consistent, since they also detected canonical changes found to be driving this mechanism among innumerous new potential candidates for acquired resistance. The integrated computational analysis were possible due to an experimental approach developed to account for molecular changes due to adaptive responses to the culturing system and the immediate addition of cetuximab. Here, we present a novel integrated analysis protocol to evaluate molecular changes measured by different high throughput techniques over a prolonged time of treatment with an FDA approved targeted therapeutic agent. The lack of *in vivo* experimentation to validate our findings was compensated by the analysis of two public datasets of HNSCC, showing that our *in vitro* findings were also present in patients’ samples. Our findings, together with Marshall et al. [47] and Marek et al. [46], are a strong evidence that FGFR1 plays a crutial role in a significant proportion of cases that are resistant to cetuximab or gefitinib. The translational implications are immense since FGFR1 inhibition can be used in combination with EGFR blockade to retard acquired resistance or overcome intrinsic resistance. It is important to mention that FGFR inhibitors are being currently evaluated by clinical trials and could soon become a potential new therapeutic option for many cancer patients [57]. Future work evaluating how these combinations impact the timing of acquired resistance are essential to determine the molecular mechanisms that shift dominant signaling pathways in cancer and thereby drive resistance.

The main limitation of the current study was the use of a single cell line model. SCC25 is intrinsically sensitive to cetuximab and from this single cell line model, we generated two groups of samples (CTX and PBS generations) over the course of 11 weeks. High throughput measurements and analysis were performed for a total of 22 samples. The collection of multiple data points in the analysis had to be accounted for when determining the number of cell lines to be included in the study. We nonetheless compared our data to gene signatures from the other isogenic HNSCC resistant model 1CC8 [10], an independent resistant model to an EGFR inhibitor in non-small cell lung cancer [38], and human tumor data from HNSCC patients prior to cetuximab treatment [42]. Besides the number of samples, we also had to take into consideration the potential batch and technical effects of broad cross-platform profiling. Nevertheless, the analysis of pretreatment HNSCC patient samples from TCGA [36] and another study [42] confirmed that our finding that *FGFR1* is up-regulated and demethylated in HNSCC and associated with acquired resistance to cetuximab is also a mechanism involved in intrinsic resistance to the targeted therapy.

The *in vitro* protocol for time course sampling developed in this study has the additional advantage of aggregating potentially heterogeneous mechanisms of resistance increasing the signal of changes in any cetuximab resistant subclone. For example, we observed demethylation and over-expression of *FGFR1* in the pooled cells, but only a single stable clone generated from the same SCC25 cell line in a previous study (CTXR10) had upregulation of *FGFR1* [31]. This finding suggests that tumor heterogeneity also plays a role in acquired resistance to targeted therapies and enables different pathways to be used to bypass the silenced target within the same tumor. Heterogeneity of SCC25 cetuximab-resistant clones has been observed previously [31]. Recent single cell RNA-sequencing data of SCC25 has shown that there is considerable transcriptional heterogeneity in this cell line prior to treatment [67]. Other cancer therapies are influenced by heterogeneity and outgrowth of resistant clones, as was observed in single cell clones isolated from the HNSCC cell line FaDu when treated with cisplatin [68]. These data and the intrinsic sensitivity of SCC25 to FGFR inhibition suggests that therapeutic resistance results from random selection of a pre-existing resistant clone. The heterogeneity in methylation profiles reflected the complexity of the resistance mechanisms that can arise from combination therapies in heterogeneous tumors. Future work extending these protocols to *in vivo* models is essential to determine the role of the microenvironment in inducing therapeutic resistance.

Developing *in vivo* models with acquired therapeutic resistance presents numerous technical challenges that must first be addressed before such time course sampling is possible [9]. Pinpointing precise molecular predictors of therapeutic resistance will facilitate the identification of unprecedented biomarkers and reveal the mechanisms by which to overcome acquired therapeutic resistance to most therapies used to treat cancer.

## CONCLUSIONS

By developing a novel bioinformatics pipeline for integrated time course analysis, we measured the changes in gene expression and DNA methylation during the progression from an intrinsic cetuximab responsive state to the acquired resistant phenotype using an *in vitro* HNSCC cell line model. Specifically, this pipeline includes (1) CoGAPS analysis of each platform independently; (2) gene selection with the PatternMarker statistics for visualization and CoGAPS gene set analysis of the CoGAPS gene profiles for pathway analysis; (3) comparisons of patterns to known phenotypes infer their relative timing; (4) anti-correlation between DNA methylation patterns and gene expression to infer epigenetically regulated genes; and (5) evaluation of PatternMarker genes and projection of the CoGAPS gene profiles to learn relevance of inferred gene signatures in new datasets. This pipeline revealed massive changes in gene expression and identified and discriminated the different patterns associated with resistance or cell culturing conditions. This analysis demonstrates that compressed sensing matrix factorization algorithms can identify gene signatures associated with the dynamics of phenotypic changes from genomics fata collected over the time course. In this case, the gene expression patterns relevant to resistance were later followed by epigenetic alterations. Our main conclusion is that using our bioinformatics approach we are able to determine that the resistant phenotype is driven by gene expression changes that would confer the cancer cells adaptive advantages to the treatment with cetuximab. Finally, the integrated analysis show that the stability of the resistant state is dependent on epigenetic changes that will make these new gene signatures heritable to expand the phenotype to the daughter cells. The bioinformatics pipeline we developed is also significant to clinical practice, since it pointed the time course of molecular changes associated with acquired cetuximab resistance and suggests that the resistant phenotype can be reversed if alternative interventions are introduced before epigenetic alterations to the genes driving acquired resistance. Most importantly the computational approach we describe here can be applied to time course studies using other tumor type models and targeted therapies.

## List of abbreviations

AKT: AKT serine/threonine kinase
ATCC: American Type Culture Collection
CoGAPS: Coordianted Gene Activity in Pattern Sets
CTX: cetuximab
CTXR: single cell cetuximab resistant clone
DNA: deoxyribonucleic acid
EGFR: Endogenous Growth Factor Receptor
FDA: Food and Drug Administration
FGFR1: Fibroblast Growth Factor Receptor 1
GEO: Gene Expression Omnibus
GSTP1: Glutathione S-Transferase pi 1
GWCoGAPS: Genome Wide Coordinated Gene Activity in Pattern Sets
HNSCC: Head and neck squamous cell carcinoma
HRAS: HRAS proto-oncogene
JHMI: Johns Hopkins Medical Institutions
LPFS: Long-progression-free survival
MAPK: Mitogen activated kinase-like protein
NSCLC: Non-small cell lung cancer
PBS: Phosphate buffered saline
PI3K: Phosphoinositide-3-kinase regulatory subunit 1
RIN: RNA integrity number
RNA: Ribonucleic acid
RNA-seq: Ribonucleic acid sequencing
rRNA: Ribosomal ribonucleic acid
SNP: Single nucleotide polymorphism
SPFS: Short-progression-free survival
STR: Short tandem repeat
TCGA: The Cancer Genome Atlas
TFAP2A: Transcription factor AP-2 alpha

## Acknowledgements

We thank JHMI Deep Sequencing & Microarray Core and SKCCC Microarray Core Facility on performing and providing advice on RNA-seq and DNA methylation hybridization arrays, respectively; S. Boca, B. Kerr, S.Floor, C. Mak, T. Ou, D. Sidransky, L. M. Weiner, F. Zamuner, K. Zambo, and members of NewPISlack for critical comments and feedback during the preparation of the manuscript.

## DECLARATIONS

### Author contributions

G.S., L.T.K, S.L., C.H.C. and E.J.F. planned, designed and wrote the manuscript with input from all authors. G.S., L.T.K., S.L., M.T., C.H.C. and E.J.F. contributed to the development of methodology. G.S., S.L. and E.J.F. performed analysis and interpretation of data (e.g., computational analysis). R.R., H.O., H.C., M.C., A.F., L.V.D., J.A., D.A.G. participated in development of methodology and provided technical and material support. R.R., L.V.D., E.I. and D.A.G. participated in review, and/or revision of the manuscript. All authors discussed the data and contributed to the manuscript preparation. C.H.C. and E.J.F. instigated and supervised the project.

## Supplementary Figures and Tables Captions

**Supplemental Figure 1 – Time course approach to induce resistance to cetuximab and measure gene expression and DNA methylation changes.** Intrinsic cetuximab sensitive HNSCC cell line SCC25 were treated with cetuximab (red) or PBS (black) for 7 days. In the eighth day, cells were collected and pooled from multiple replicate cultures to provide adequate amounts for total RNA isolation for RNA-seq, genomic DNA isolation for DNA methylation array, proliferation assay (flow), for storage (frozen) and to be plated again to continue treatment until resistance to cetuximab developed. Each collection point was called a generation (from CTX-G0 to CTX-G11).

**Supplemental Figure 2 –** Colony formation assay in matrigel for anchorage-independent growth confirmed acquired cetuximab resistance of CTX-G10 (red) relative to the parental cell line (CTX-G0, black) at different concentrations of cetuximab (0nM, 10nM, 100nM and 1000nM).

**Supplemental Figure 3 –** Heatmap and hierarchical clustering of gene expression values in 11 generations of SCC25 cells treated with PBS as control (black columns) and with 100nM of cetuximab (red columns) to acquire resistance.

**Supplemental Figure 4 – A.** Heatmap of gene expression values in 11 generations of SCC25 cells treated with 100nM of cetuximab (red columns) to acquire resistance and with PBS as control (black columns). Genes selected for visualization were associated with cetuximab resistance from previous gene expression studies comparing sensitive and resistant cells without regard for timing. These studies provided three gene sets, colored along rows of the heatmap. **B.** Average of z-score gene expression values for genes in each of the resistance signatures over generations of PBS control (black lines) or treatment with 100nM of cetuximab (red lines).

**Supplemental Figure 5 –** Expected gene expression values for genes in each CoGAPS pattern inferred from gene expression data over generations of PBS control (black lines) or treatment with 100nM of cetuximab (red lines). Patterns included a pattern reflecting technical artifacts between untreated controls at time 0 and subsequent generations (pattern 4) and a flat pattern for highly expressed genes (pattern 5), excluded from analysis in main figures. Heatmap of gene expression values for PatternMarker genes identified for all of these patterns. Rows were colored according to which CoGAPS pattern the PatternMarker statistic assigned each gene, and sorted by the PatternMarker statistic.

**Supplemental Figure 6 –** Heatmap of gene set analysis scores for targets of transcription factors in the EGFR network, targets of the AP-2alpha transcription factors associated with cetuximab response, and cetuximab resistance signatures in CoGAPS patterns. A score of 100 indicated upregulation of the targets with a p-value of 0 and −100 downregulation with p-values of 0. Matrix elements with a star indicated p-values below 0.05 for either up or down-regulation of the gene set. Gene expression heatmap was colored on a red-green scale where as the gene set statistics heatmap was colored on a blue-red scale, with values indicated in the respective color keys.

**Supplemental Figure 7 – A.** Heatmap of Pearson correlation coefficients between CoGAPS gene expression and DNA methylation patterns. Row colors for expression patterns match the colors for patterns in Figure 2,3. The column colors for methylation patterns are selected to match the color of the corresponding expression pattern with maximum anti-correlation. **B**. As in **A** for CoGAPS gene weights (meta-pathway values) corresponding to patterns in DNA methylation (columns) and gene expression (rows).

**Supplemental Figure 8 –** Heatmap of gene expression values for 11 generations of SCC25 cells treated with PBS as control (black columns labeled PBS) and with 100nM of cetuximab (red columns labeled cetuximab) to acquire resistance and gene expression data from independent, stable cetuximab resistant clones in absence of cetuximab treatment (CTX resistant clones).

**Supplemental Figure 9 –** Heatmap of DNA methylation values for 11 generations of SCC25 cells treated with PBS as control (black columns labeled PBS) and with 100nM of cetuximab (red columns labeled cetuximab) to acquire resistance and gene expression data from independent, stable cetuximab resistant clones in absence of cetuximab treatment (CTX resistant clones).

**Supplemental Figure 10 –** Brightfield microscopy images of the representative clones’ morphology above with corresponding clone specific CoGAPS patterns of DNA methylation above.

**Supplemental Figure 11 –** Epigenetically regulated pattern marker genes associated with resistance having significant anti-correlation between gene expression and DNA methylation in the cetuximab single cell resistant clones.

**Supplemental Figure 12 –** Cell proliferation assay using AlamarBlue (Invitrogen, Carlsbad, CA) to compare proliferation rates under different concentrations of cetuximab in the resistant single cell clones (CTXR4, 7, 10 and 11) and the parental SCC25 cell line to confirm resistance when treated with different concentrations of cetuximab.

**Supplemental Figure 13 –** Heatmap of FGF family genes methylation **(A)** and expression **(B)** in time course data for treated and control samples.

**Supplemental Figure 14 –** Bar plot of FGFR1 expression in human tumor samples pre (black) and post (red) treatment with cetuximab.

**Supplemental Table 1 –** PatternMarker genes for the expression and methylation CoGAPS patterns and list of genes with significant anti-correlation between gene expression and methylation.

**Supplemental Table 2 –** P-values of MSigDB hallmark pathways for CoGAPS Expression patterns calculated with the permutation-based statistic in calcCoGAPSStat.

**Supplemental Table 3 –** P-values of MSigDB hallmark pathways for CoGAPS DNA Methylation patterns calculated with the permutation-based statistic in calcCoGAPSStat.

